# Metabolic cost calculations of gait using musculoskeletal energy models, a comparison study

**DOI:** 10.1101/588590

**Authors:** Anne D. Koelewijn, Dieter Heinrich, Antonie J. van den Bogert

**Affiliations:** Parker Hannifin Laboratory for Human Motion and Control, Department of Mechanical Engineering, Cleveland State University, Cleveland, Ohio, USA; Biorobotics Laboratory, Institute of Bioengineering, École Polytechnique Fédérale de Lausanne, Lausanne, Switzerland; Department of Sport Science, University of Innsbruck, Innsbruck, Austria

## Abstract

This paper compares predictions of metabolic energy expenditure in gait using seven metabolic energy expenditure models to assess their correlation with experimental data. Ground reaction forces, marker data, and pulmonary gas exchange data were recorded for six walking trials at combinations of two speeds, 0.8 m/s and 1.3 m/s, and three inclines, −8% (downhill), level, and 8% (uphill). The metabolic cost, calculated with the metabolic energy models was compared to the metabolic cost from the pulmonary gas exchange rates. A repeated measures correlation showed that all models correlated well with experimental data, with correlations of at least 0.9. The model by Bhargava et al. [7] and the model by Lichtwark and Wilson [21] had the highest correlation, 0.96. The model by Margaria [23] predicted the increase in metabolic cost following a change in dynamics best in absolute terms.

## Introduction

Humans prefer to walk in energetically optimal ways. Walking speed [1], the ratio between step length and frequency [2], step width [3] and vertical movement of the center of mass [4, 5] are chosen to minimize energy expenditure. Whole-body energy expenditure can be measured using direct calorimetry, by measuring the heat production in the body, or indirect calorimetry, by measuring the volume of oxygen inspired and carbon dioxide expired [6]. However, these measurements are not always available, but an accurate prediction of energy expenditure is often desired.

Instead, musculoskeletal modeling can be used to simulate human gait and to calculate the energy expenditure based on a metabolic energy expenditure model (e.g., [7–10]). The metabolic cost of walking is the energy expended by the human body to move a certain distance. The metabolic cost can be calculated using variables that are studied in gait analysis, such as joint moments, joint power, or muscle forces, lengths and activations [11]. With a metabolic energy expenditure model, the energy expenditure can be calculated for gait experiments that did not take metabolic energy expenditure measurements, or for predictive gait simulations, where no experimental data is available. Recently, these simulations have been used to analyze ‘what-if scenarios’ such as the effect of an intervention such as a prosthesis [12], an exoskeleton [13], ankle foot orthosis [14], additional weight [15], loaded and inclined walking [16], or changing the gait pattern to minimize knee reaction force [17] on gait. Energy cost is an important variable in gait, and should be considered whenever a clinical intervention is studied or designed.

In literature several energy models were suggested. The Huxley crossbridge model [18] finds both the muscle force and the energy expenditure of a muscle, but requires up to 18 states [19]. Instead, Hill-type muscle models [20] are typically used to simulate muscles, but these do not output metabolic energy expenditure. Therefore, several metabolic energy expenditure models have been proposed that calculate the energy expenditure during walking based on Hill-type muscles [7–10, 21, 22], based on muscle efficiency [23], or based on joint angles and moments [24]. An additional model is based on another, similar muscle model [25]. An additional advantage of metabolic energy expenditure models is their ability to calculate the energy expenditure of single muscles [22], or joints [26] and provide more information than measurements of pulmonary gas exchange, which only calculated on the whole body energy expenditure.

So far, these models have only been compared and used on level walking studies and self-selected speed (e.g. [7, 9, 26]). However, it is important to know how well these models can represent changes in energy cost due to altered control, such as a change in walking speed, or environment. If the representation is accurate, these models can be used to assess the energetic effect of an intervention.

Consequently, we aimed to compare metabolic cost calculated with the different models to metabolic cost measured with indirect calorimetry on walking trials with different speeds and slopes. These were used as test cases for intervention studies, since it requires a similar change in dynamics, while there is some information in literature of the effect of these dynamics changes to energy expenditure. It is known that in downhill walking, knee extensor activity increases [27, 28] while metabolic cost decreases [23], that in uphill walking metabolic cost increases [23], and that between 2 and 5 km/h, metabolic cost is independent of speed [29]. Therefore, a secondary goal is to see if the metabolic cost calculated with metabolic energy expenditure models has a similar effect with a change in slope or speed.

## Methods

### Subjects and experiment

Twelve healthy participants (6 female, 6 male, mean *±* SD age 24 *±* 5 years, weight 70 *±* 12 kg, and height 173 *±* 8 cm) were recruited using flyers at Cleveland State University and word-of-mouth. They performed the experiment after providing informed consent. The experimental protocol was approved by the institutional review board of Cleveland State University (IRB-FY2017-286). First, the subjects stood on the treadmill for three minutes to determine their resting metabolic rate. They performed six walking trials of seven minutes in random order, three at 0.8 m/s and three at 1.3 m/s. For each speed, there were three different inclines: level walking, downhill walking with a negative incline of 8%, and uphill walking with a positive incline of 8%. Pulmonary gas exchange rates were measured with the COSMED K4b2 system (COSMED, Italy). An instrumented treadmill with two six degree of freedom force plates (R-Mill, Forcelink, Culemborg, the Netherlands) was used to measure the ground reaction forces. A motion capture system with 10 Osprey cameras and Cortex software (Motion Analysis, Santa Rosa, CA) was used to record 27 markers, given the markerset in S1 Fig and S1 Table. The raw and processed experimental data are published as a dataset in [30].

### Metabolic energy models

Seven metabolic energy models were selected for the current study: models BHAR04 [7], HOUD06 [10], UMBE03 [9], LICH05 [21], MINE97 [8], MARG68 [23], and KIMR15 [24]. Six models use muscle states (contractile element length, activation, stimulation) to determine the energy rate of the individual muscles. Model KIMR15 calculates the energy rate for each joint instead of each muscle, using the angular velocity and joint moment.

The calculated metabolic cost of walking, *C*_*calc*_, is determined in J/kg/m as follows:

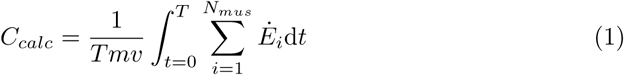

where *T* denotes the motion duration, *m* the participant’s mass, *v* the speed, *N*_*mus*_ the number of muscles, and *Ė*_*i*_ the energy rate of muscle *i* in W.

Models BHAR04, HOUD06, UMBE03, and LICH05 calculate the energy rate as a function of work rate, 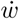, and heat rates, due to activation, 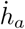, maintenance of contraction, and 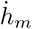, and muscle shortening and lengthening, 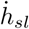 [7, 9, 10, 21]:

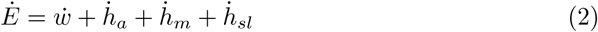

The implementation of these models is detailed in S1 File.

Model MINE97 determines the energy rate for each muscle incorporating an empirical function of the ratio between the contractile element velocity, *v*_*CE*_ and the maximum contractile element velocity, *v*_*CE*(*max*)_ [8]:

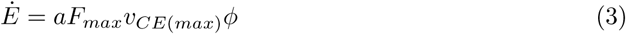

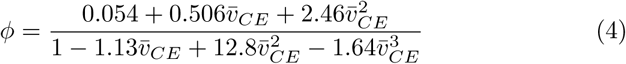

where *a* is the muscle activation, *F*_*max*_ the maximum isometric force, and 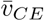 is the ratio of the contractile element velocity to the maximum contractile element velocity.

Model MARG68 is based on the observation that muscles are 25% efficient when shortening, and 120% efficient when lengthening [23]:

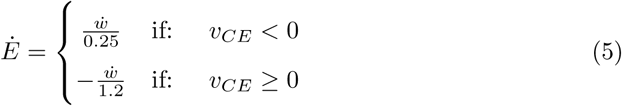

Model KIMR15 does not use muscle states, but calculates the metabolic rate on the joint level, using the joint moments and angular velocities. The metabolic rate is still the sum of the heat rate and the work [24]:

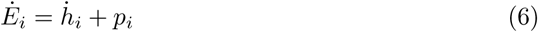

where the power, *p*_*i*_, at joint *i* is the product of the joint moment *M* and angular velocity 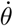 [24]:

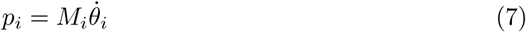

The heat rate is determined as follows [24]:

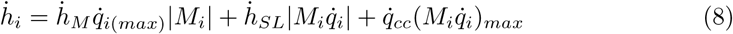

where 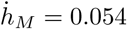 is the heat rate for activation and maintenance, 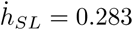 is the shortening-lengthening heat rate for positive power, and 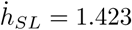 is the shortening-lengthening heat rate for negative power, and 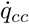 is the cocontraction heat rate. The subscript *max* indicates the maximum over the gait cycle [24].

### Kinetic and Kinematic Data Processing

A two step approach was used calculate the joint angles, moments, and muscle states and inputs necessary to determine the metabolic cost of walking level, uphill and downhill using the metabolic energy models.

In the first step, the joint angles and moments were determined from marker and ground reaction force data. The data was filtered backwards and forwards with a second order Butterworth filter with a cut-off frequency of 6 Hz. Velocities and accelerations were calculated using finite differences from marker positions. The data was split into gait cycles and resampled to 100 data points per gait cycle.

The joint angles were determined using the orientation from the proximal to the distal marker on the body segment. For example, for the tibia these were the knee and ankle markers (RLEK/LLEK and RLM/LLM, see S1 Table and S1 Fig). The joint moments were determined from the marker data and the ground reaction forces using Winter’s method [31]. The joint angles, moments, and ground reaction forces were averaged over all left and right gait cycles to find one average gait cycle.

In the second step, the muscle states (activation and contractile element length) and stimulations were determined using the dynamic optimization method introduced in [32] such that the joint moments generated by the muscles matched the moments found using Winter’s method. Fig 1 shows the eight Hill-type muscles that were modeled in each leg. These muscles were modeled as three element Hill-type muscles with quadratic springs for the parallel and series elastic element. The contractile element had activation dynamics, a force-length relationship, and a force-velocity relationship [15]. A full description of the muscle model and the muscle parameters are listed in S2 File.

**Fig 1.**
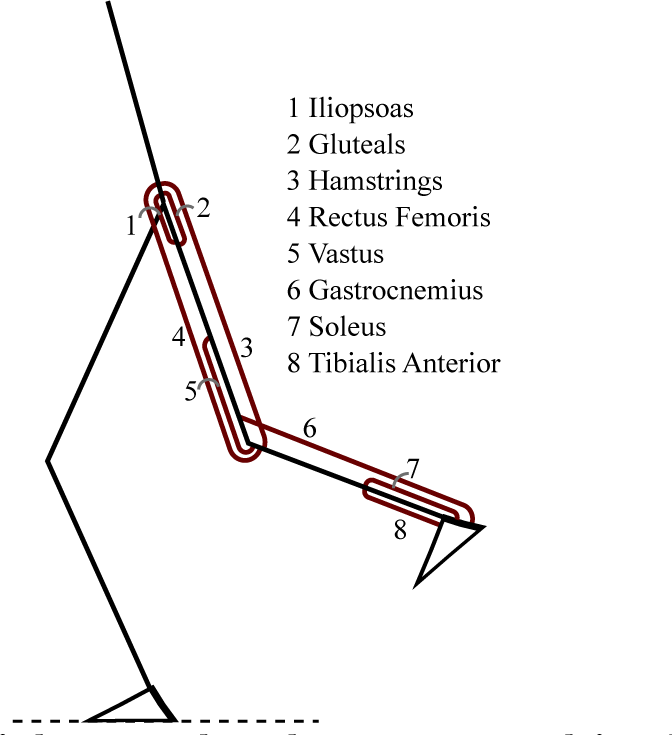
Schematic of the eight muscles that were used in the current study.

The stimulations *u*(*t*), activations *a*(*t*), and contractile element lengths *l*_*CE*_(*t*) were found by solving the following dynamic optimization problem:

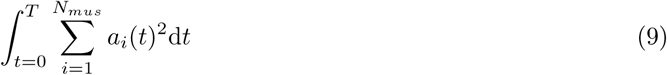

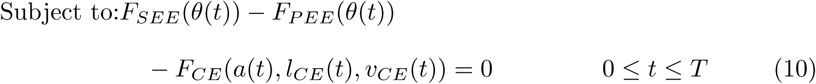

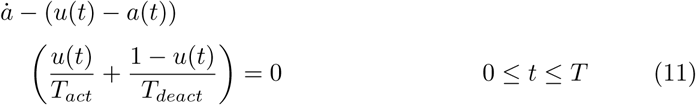

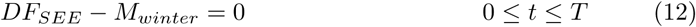

where *F*_*SEE*_, *F*_*PEE*_ and *F*_*CE*_ denote the series elastic, parallel elastic and contractile element force, *θ* the joint angles, *v*_*CE*_ the contractile element velocity, the first derivative of the contractile element length. *T*_*act*_ is the activation time constant, *T*_*deact*_ is the deactivation time constant, *D* denotes a matrix of muscle moment arms, and *M*_*winter*_ the moments that were calculated previously. Periodic boundary conditions were used: *u*(*T*) = *u*(0), *a*(*T*) = *a*(0), and *l*_*CE*_(*T*) = *l*_*CE*_(0).

This dynamic optimization problem was solved using direct collocation, with 100 nodes for the gait cycle and a Backward Euler formulation. IPOPT 3.11.0 was used to solve the optimization problem [33]. Muscle force patterns of level walking at 1.3 m/s were compared to electromyography (EMG) data by Winter and Yack [34]. The EMG data was digitized from the paper, and averaged when more than one muscle from a muscle group was recorded (e.g. medial and lateral hamstrings). Finally, the muscle state trajectories and stimulations, or the joint angular velocities and moments were inserted in the seven metabolic energy models to find the calculated metabolic cost.

### Pulmonary Data Processing

The measured metabolic cost was derived from the pulmonary gas exchange data using indirect calorimetry. The first 30 seconds of the resting trial, and the first three minutes of each walking trial were disregarded. The rate of oxygen consumption, 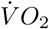 in mL/min/kg, and respiratory quotient, *R* were averaged over time. The metabolic rate in W/kg was determined as follows for the resting and walking trials [35, 36]:

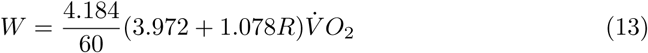

The resting trial was subtracted from each walking trial. The metabolic rate was divided by walking speed to find the measured metabolic cost in J/kg/m:

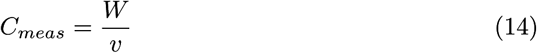

### Analysis

First, the implementation of the metabolic energy models was verified and compared to Miller [26]. To do so, the metabolic rate, *Ė*, was determined for the soleus for three speeds (shortening at 1 *l*_*ce*_/s, isometric and lengthening at 1 *l*_*ce*_/s), and five activation levels (0.05, 0.25, 0.5, 0.75 and 1). The stimulation was assumed to be the same as the activation. The stimulation time, used in models BHAR04 and LICH05, was set to 1.

After the verification of the metabolic energy models, the metabolic cost was calculated for every subject and trial (Eq. 1) and compared to the measured metabolic cost (Eq. 2). Additionally, the calculated metabolic cost was determined for all joints individually. The metabolic cost of biarticular muscles was split between the joints on which they act using the ratios of the moment arms. The difference between the calculated and measured metabolic cost, averaged over all subjects, is also presented.

Finally, we investigated the ability of the metabolic energy models to predict a change in energy following a change in dynamics - due to altered walking speed and/or incline. Specifically, a repeated measures correlation [37] was used to determine how well the calculated metabolic cost correlated with the measured metabolic cost for every trial. A repeated measures analysis can account for the fact that more than one data point (i.e. different speeds and slopes) is available for each subject, which creates a dependency between the data points. A repeated measures analysis accounts for multiple data points for each subject by assuming a model with the same slope for each subject, but a different intercept. The analysis with a fixed slope was chosen since the slope could then be applied to predict changes in metabolic cost for new subjects. The intercept was not fixed since it is not relevant when predicting a change in metabolic energy.

## Results

### Verification of Metabolic Energy Models

Fig 2 shows the metabolic power of the soleus muscle for several activation levels and a shortening and lengthening velocity of 1 *l*_*CE(OPT)*_/s, where *l*_*CE(OPT)*_ is the optimal contractile element length, and an isometric condition. The metabolic rate for shortening is more than twice as high for model MARG68 than for all other models. In the isometric condition model MINE97 has the largest metabolic rate, while model MARG68 has zero metabolic rate, since no work is done at zero speed. The metabolic rate is most different between the models during lengthening, where models MARG68 and MINE97 had a positive metabolic rate, and the other models had a negative rate.

**Fig 2.**
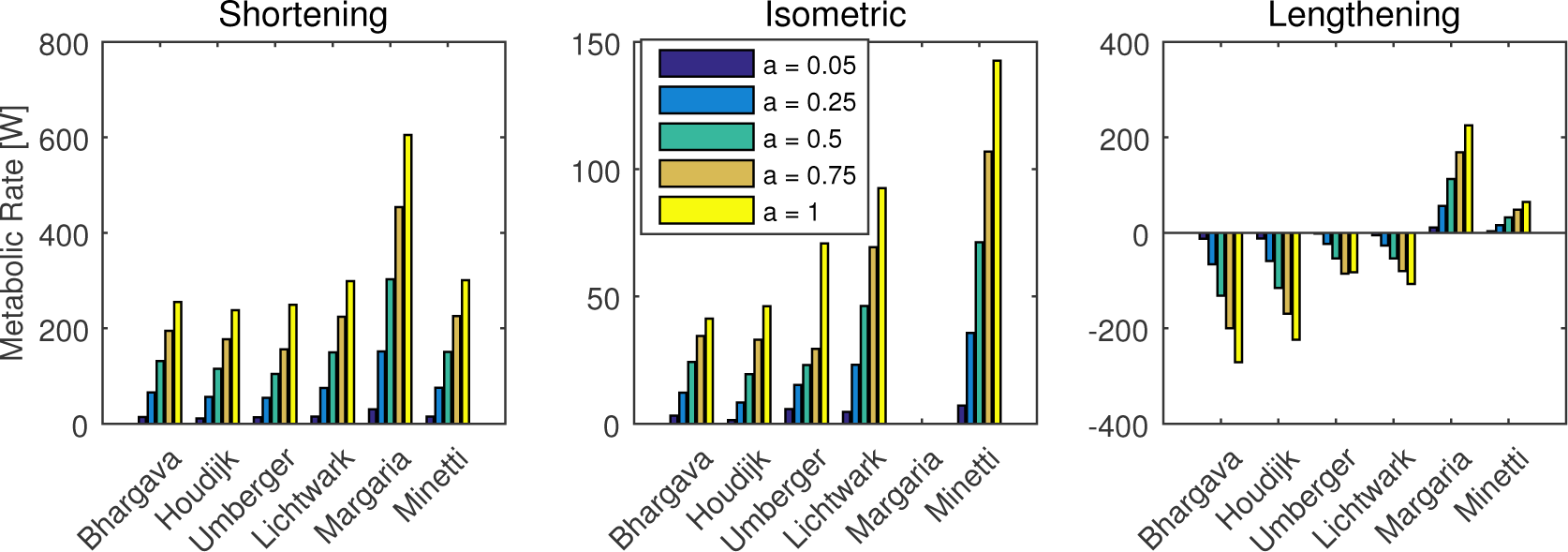
Metabolic rate of soleus, calculated with the metabolic energy models for different conditions. Metabolic rate as calculated by the metabolic energy expenditure models that are based on individual muscles of the soleus muscle at four activation levels, for an isometric condition, and a shortening and lengthening velocity of one optimal fiber length per second.

### Joint Kinetics and Kinematics

Fig 3 shows the measured ground reaction forces, and the calculated joint angles, joint moments, and muscle forces for all trials at 1.3 m/s. The black dashed line in the muscle force graph shows EMG data for comparison [34], scaled to the maximum muscle force. The muscle force pattern matches the EMG data. In the gastrocnemius, the EMG activation increases earlier in the gait cycle than the muscle force, while in the rectus femoris, there is an extra burst in the muscle force at about 60-80% of the gait cycle, and in the tibialis anterior, the activity of the EMG is generally higher than the muscle force.

**Fig 3.**
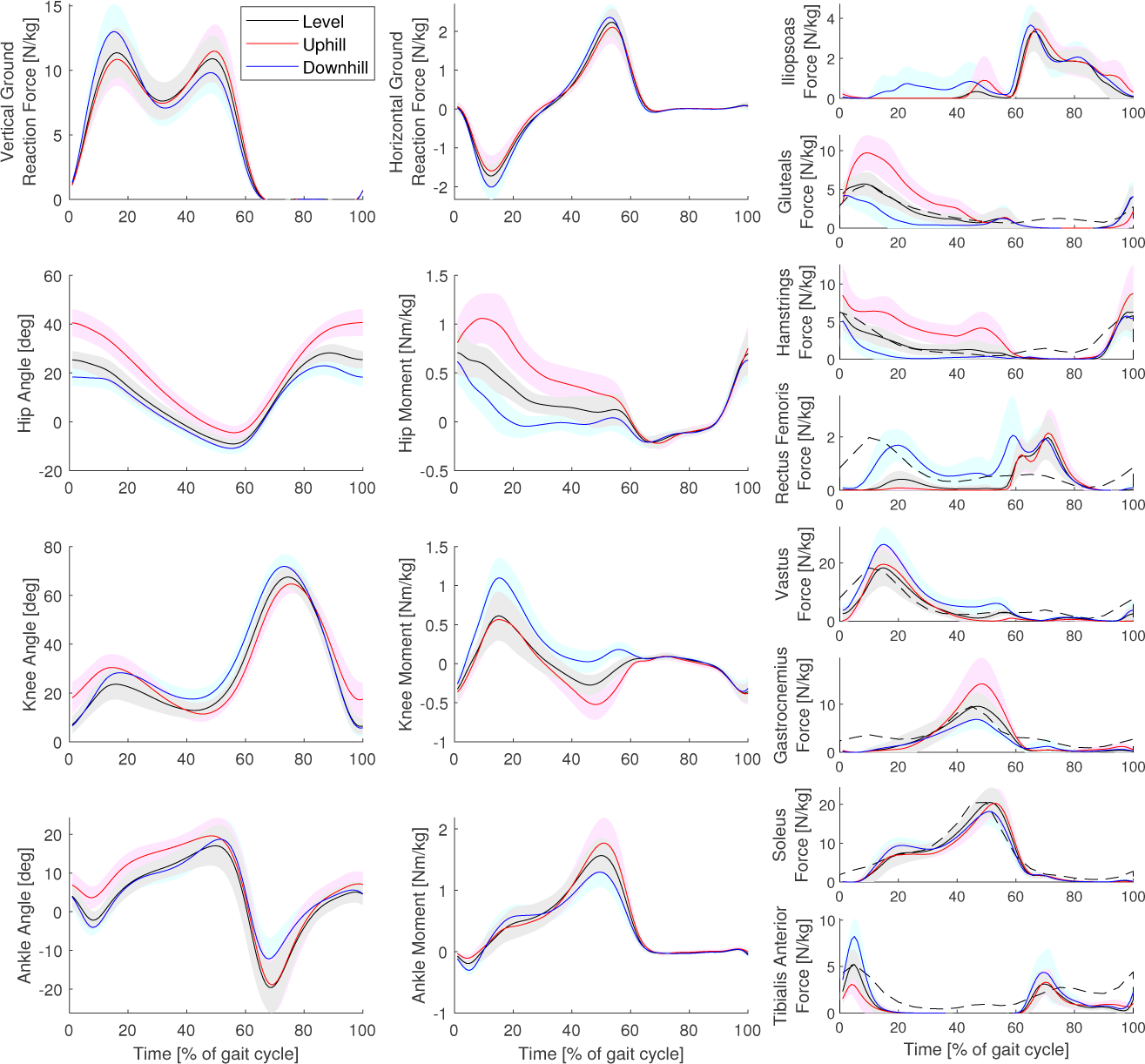
Kinetics and kinematics at the normal speed. Average ground reaction forces, joint angles, joint moments, and muscle forces for all trials at the normal speed (1.3 m/s). The shaded area denotes one standard deviation. The dashed line shows EMG data of level walking for comparison [34]. The graphs use Winter’s sign convention, where flexion angles and extension moments are positive for hip and knee. Dorsiflexion angle and plantarflexion moment are positive for the ankle.

Fig 4 shows the measured ground reaction forces, and the calculated joint angles, joint moments, and muscle forces for all trials at 0.8 m/s. The pattern of the downhill, level, and uphill trials in the joint angles, moments, and ground reaction forces were very similar between the two speeds. The muscle forces were lower at 0.8 m/s, but showed similar trends as in 1.3 m/s, except for the vastus and soleus.

**Fig 4.**
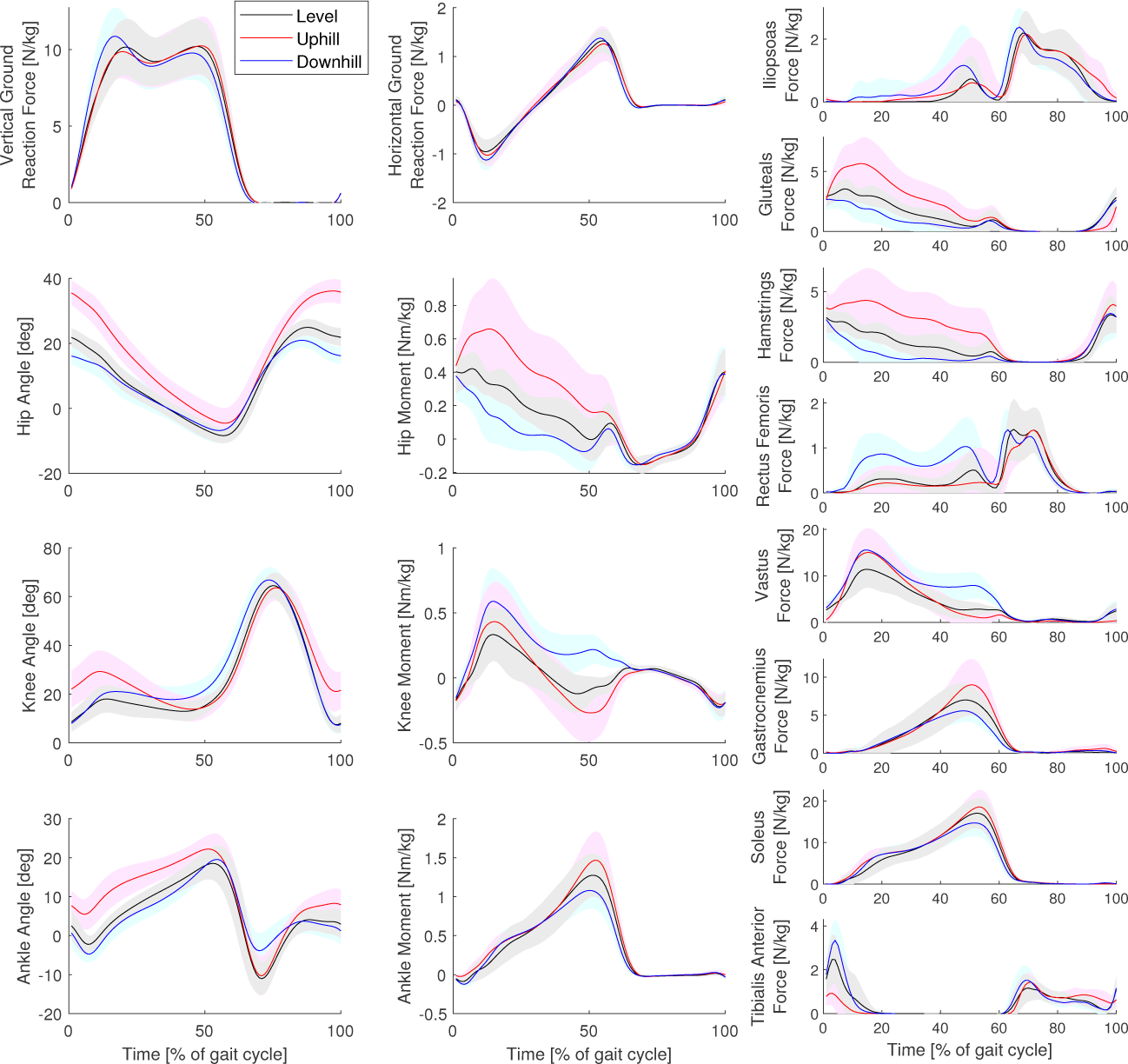
Kinetics and kinematics for the slow speed. Average ground reaction forces, joint angles, joint moments, and muscle forces for all trials at the slow speed (0.8 m/s). The shaded area denotes on standard deviation. The graphs use Winter’s sign convention, where flexion angles and extension moments are positive for hip and knee. Dorsiflexion angle and plantarflexion moment are positive for the ankle.

### Comparison of Calculated and Measured Metabolic Cost

Fig 5 shows the calculated metabolic cost for each model, separated for the hip, knee and ankle joints. Biarticular muscles were added by ratio of the moment arm, similar to [26]. The mean calculated metabolic cost was lowest for model HOUD06 and highest for model MARG68, ranging from (mean *±* SD): 0.31 *±* 0.14 J/kg/m to 3.0 *±* 0.36 J/kg/m for the downhill trials, from 1.1 *±* 0.19 J/kg/m to 4.3 *±* 0.37 J/kg/m for the level trials and from 2.2 *±* 0.33 J/kg/m to 6.3 *±* 0.40 J/kg/m for the uphill trials. The mean measured metabolic cost ranged from 2.0 *±* 0.40 J/kg/m for the downhill trial at 1.3 m/s to 5.9 *±* 0.34 J/kg/m for the uphill trial at 1.3 m/s (see bottom right graph). One measurement was missing for the downhill trial at 1.3 m/s due to a malfunction of the K4b2 system.

**Fig 5.**
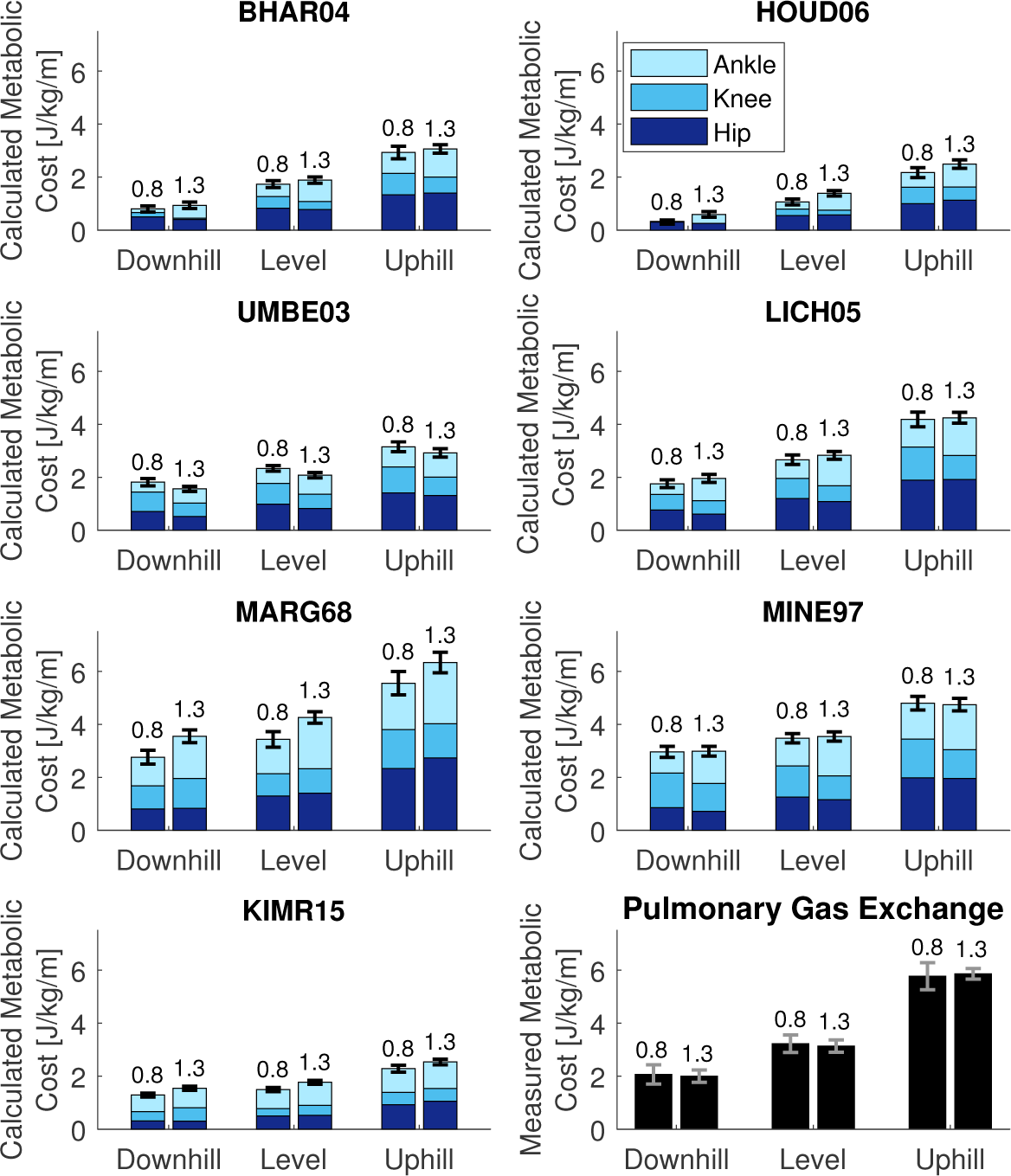
Calculated and measured metabolic cost for all trials and models. Calculated metabolic cost for all speeds and inclines, for each model, separately for the joints. The number above the bar indicates the walking speed. Biarticular muscles were added by ratio of the moment arm, similar to [26]

Comparing the different inclines, the calculated metabolic cost increased with incline for all models. The energy expenditure at the hip increased most from downhill to level to uphill walking. The metabolic cost calculated with model BHAR04 and model HOUD06 also increased at the knee and ankle joint with an increasing slope.

The effect of speed differed among the models. With model BHAR04, model HOUD06, model UMBE03, model LICH05, and model MINE97, the metabolic cost shifts from the knee to the ankle when the speed increases. For model UMBE03, model LICH05, and model MINE97, the summed metabolic cost of the knee and ankle is the same for both speeds. With model MARG68, there is an increase in metabolic cost at the ankle with speed, while for model KIMR15 there is an increase in the hip and ankle.

Table 1 shows the root mean square (RMS) error between the measured metabolic cost and the calculated metabolic cost. Model LICH05 and UMBE03, and model KIMR15 at 1.3 m/s produced the lowest RMS error in the downhill trials (between 0.54 and 0.71 J/kg/m), while model HOUD06 produced the largest error at 0.8 m/s (1.86 J/kg/m). Model MINE97 produced the lowest RMS error in the level trials (0.59 and 0.66 J/kg/m), while model HOUD06 and KIMR15 at 0.8 m/s produced the highest errors (at least 1.78 J/kg/m). Model MARG68 produced the lowest RMS error in the uphill trials (0.81 J/kg/m), while model KIMR15 and model HOUD06 produced the highest error (at least 3.34 J/kg/m). The RMS errors in the uphill trials were higher than in the level and downhill trials for all models except MARG68. In the level and downhill trials, only model HOUD06 produced an RMS error larger than 2.0 J/kg/m, at 0.8 m/s in the level trial.

**Table 1.** Root mean square (RMS) error between the calculated and measured metabolic cost in J/kg/m for all models and all trials.

| Model | Downhill |  | Level |  | Uphill |  |
| --- | --- | --- | --- | --- | --- | --- |
|  | 0.8 m/s | 1.3 m/s | 0.8 m/s | 1.3 m/s | 0.8 m/s | 1.3 m/s |
| BHAR04 | 1.41 | 1.08 | 1.59 | 1.31 | 2.92 | 2.82 |
| HOUD06 | 1.86 | 1.39 | 2.23 | 1.78 | 3.68 | 3.39 |
| UMBE03 | 0.69 | 0.53 | 1.05 | 1.11 | 2.73 | 2.96 |
| LICH05 | 0.71 | 0.33 | 0.83 | 0.48 | 1.76 | 1.67 |
| MARG68 | 0.93 | 1.46 | 0.78 | 1.19 | 0.90 | 0.81 |
| MINE97 | 1.12 | 0.97 | 0.66 | 0.59 | 1.26 | 1.21 |
| KIMR15 | 0.98 | 0.54 | 1.82 | 1.43 | 3.59 | 3.34 |

Fig 6 shows the linear regression model that was fitted during the correlation analysis for each metabolic energy model. Table 2 shows the correlation coefficients of all models. All models correlate with a coefficient of at least 0.9. The highest correlation coefficient was 0.96 for model BHAR04 and model LICH05. The lowest correlation coefficient was 0.90 for model KIMR15 and model MARG68. The slope of the regression model for model MARG68 (1.13) was closest to unity, while the slope for model KIMR15 (3.20) was furthest away from unity.

**Table 2.** Results of repeated measures correlation. Repeated measures correlation coefficient *r*_*rm*_ [37] with 95% confidence interval (CI) and slope of the repeated measures model.

| Model | $r_{rm}$ | (95 % CI) | Slope |
| --- | --- | --- | --- |
| BHAR04 | 0.96 | (0.93 - 0.97) | 1.75 |
| HOUD06 | 0.94 | (0.90 - 0.96) | 1.92 |
| UMBE03 | 0.93 | (0.89 - 0.96) | 2.62 |
| LICH05 | 0.96 | (0.93 - 0.97) | 1.56 |
| MARG68 | 0.90 | (0.84 - 0.94) | 1.13 |
| MINE97 | 0.95 | (0.92 - 0.97) | 1.98 |
| KIMR15 | 0.91 | (0.85 - 0.95) | 3.20 |

**Fig 6.**
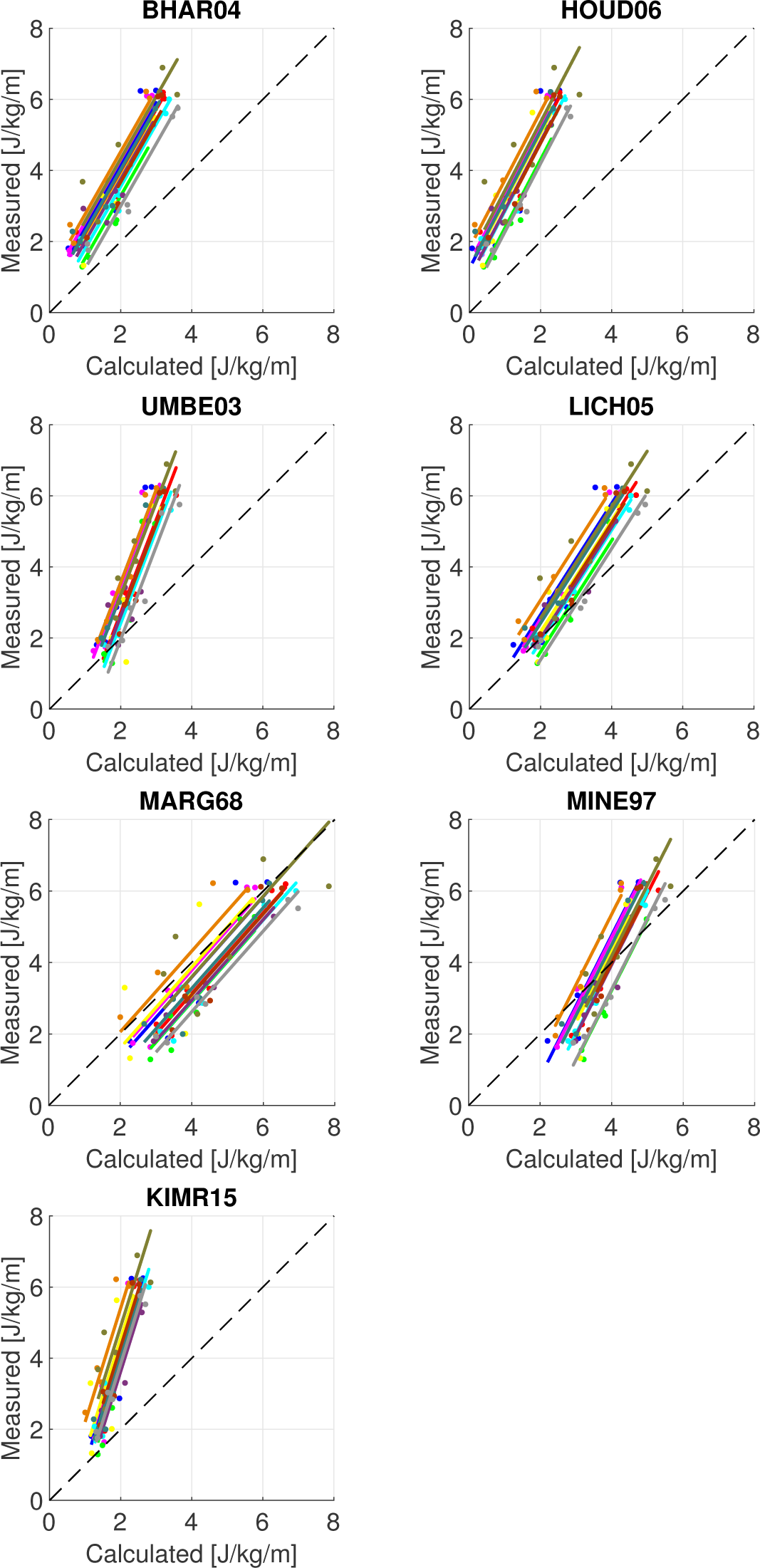
Correlation graphs for all models. Correlation graphs between calculated and measured metabolic costs for each model. The lines show the regression model that was fitted by the repeated measured correlation.

## Discussion

The goal of the current study was to compare metabolic cost calculated with seven metabolic models to metabolic cost measured with indirect calorimetry on walking trials with different speeds and slopes. All models correlated with a coefficient of at least 0.9, while model BHAR04 and model LICH05 correlated best with the experimental data (see Table 2). The regression model of model MARG68 had a slope closest to unity, which indicates that model MARG68 calculates absolute differences between two trials most accurately. All models had a slope larger than one, meaning that the trials with higher metabolic demands were underestimated to a larger extent than those with a smaller metabolic demand, which is also visible in Table 1, where the RMS error of the uphill trials was consistently higher than the downhill and level trials. All models predicted a lower metabolic cost in the downhill trials than in the level trials, despite a larger force in the knee extensors (rectus femoris throughout stance and vastus during late stance), similar to observed in previous studies [23, 27, 28]. Model UMBE03 predicted a slightly lower metabolic cost for the larger speed, while model LICH05 and MINE97 predicted a similar cost between speeds, which is similar to the measured metabolic cost and [29], and the other models predicted a slightly higher metabolic cost for the larger speed. The high correlation confirms Hicks et al. [38], who mention that a sagittal plane model should be sufficiently accurate for walking, since walking occurs almost entirely in the sagittal plane. Correlation coefficients were also high for all subjects individually.

Most models, except MINE97 and MARG68, generally underestimate the metabolic cost. A possible reason is the approach used to find the muscle activations by minimizing muscular effort. Then, muscular co-contraction was disregarded. However, the muscle forces matched EMG patterns (see Fig 3), which supports that the muscle activation pattern is accurate, but the quality of the activation level cannot be assessed. Additionally, the model used in the current study was a two-dimensional model of the trunk and lower leg, meaning that metabolic cost due to lateral and rotational motion, as well as arm swing, were not taken into account, which could cause an underestimation of metabolic cost. Finally, the calculated metabolic cost could be lower since negative work was subtracted in the current study. However, the high correlation indicates that the mechanical analysis still yielded a correct prediction of increases or decreases in metabolic cost, which was the main aim of the current study.

The calculated metabolic cost in the current study is lower than calculated by Bhargava et al. [7] at 1.36 m/s (1.9 *±* 0.20 J/kg/m vs. 4.3 J/kg/m) and Umberger et al. [9] at 1.2 m/s: (2.1 *±* 0.18 J/kg/m versus 3.7 J/kg/m). Both models included a resting metabolic rate, and used a three dimensional model [7, 9], while the current study used a sagittal plane model with eight muscles and no resting metabolic rate. When accounting for a resting metabolic rate of 1.2 W/kg (see [9]), the calculated metabolic cost was still lower in the current study. Both studies minimized metabolic cost, and therefore a lower metabolic cost was expected in these works than in the current study. However, more recently, the metabolic models of Bhargava et al. and Umberger et al. were used as objectives in predictive simulations [39, 40]. In these recent studies, the metabolic cost of the simulations was lower than in the current work, and much lower than reported by Bhargava et al. [7] and Umberger et al. [9]. Therefore, the underestimation was not unexpected. While the dimensionality of the model played a role, the difference in metabolic cost between the original and more recent work could also be caused by the limited number of parameters that were used for the muscle stimulation or due to differences in kinetics or kinematics, which were not reported by either [7, 9]. Nonetheless, the patterns between different conditions were similar to previous work. Model UMBE03 underestimated the increase in metabolic cost from level to uphill, and the decrease in metabolic cost from level to downhill. Dembia et al. [13] found a similar result using model UMBE03 to predict the increase in metabolic cost from unloaded walking to loaded walking with 38 kg on the torso.

The simplest model, MARG68, had a slope closest to 1, meaning that this model was best able to predict the absolute change in metabolic cost following the changes in the dynamics. Model MARG68 was based on observations on slope walking, which was the change in dynamics used in the current study. Additionally, model MARG68 is based only on work, which is the main factor that differs with a change in slope. It was also observed that the simple models (MARG68 and MINE97) yielded some of the most accurate results. However, it was found that the absolute performance of the metabolic energy models was dependent on different factors, such as the equation that was used to determine the measured metabolic cost and the resting metabolic cost that was used, both of which are discussed later in this section. Therefore, the absolute performance of the metabolic energy models should be investigated further.

The handling of muscular energy expenditure during lengthening is still debated [26]. During lengthening, the energy rate can be negative, which is physically impossible. However, the negative work should be subtracted from the metabolic cost in models BHAR04, HOUD06, UMBE03, and LICH05 [7, 9, 10, 21]. We aimed to see if predictions improved without subtracting negative work, which is physically more sensible. When negative work was not subtracted, the calculated metabolic cost increased. The RMS error decreased, except for model LICH05 in the downhill trials. The correlation coefficient remained 0.96 for model BHAR04, but decreased for all other models, 0.01 for model LICH04, 0.02 for model HOUD06, and 0.03 for model UMBE03. The lenghtening heat rate coefficient was updated in model UMBE03 according to [41], so it is interesting that the difference was largest for model UMBE03.

The models assumed an equal maximum isometric force, and thus equal muscle mass, for all participants, even though their body masses were different. A sensitivity analysis was done to determine whether this could have affected the results. The maximum isometric force was increased and decreased 10% from the nominal value for all participants, and personalized using the ratio of the subject’s weight to the average, meaning that if the weight was 20% above average, the force was multiplied with 1.2. These changes only affected the correlation and slope very slightly. When the force was personalized, the correlation of model UMBE03 increased 0.01. The largest change in the slope was 0.06 for model UMBE03 when the force was personalized. Additionally, the muscle stress was doubled such that the muscle weight (see S2 File) was equal to about 50% of the leg weight [26], which caused the correlation of model UMBE03 to increase by 0.03, and the correlation of BHAR04 (0.02) and HOUD06 (0.01) to decrease. This was the only case that was studied where the correlations of model BHAR04 and LICH05 were not the highest, but model UMBE03 and model LICH05 performed best.

The kinematic and kinetic data are similar to previous experimental studies of sloped walking [28, 42]. The trend of the muscle forces with the slope was similar to [43] for the gluteals, hamstrings, rectus femoris and gastrocnemius. Alexander and Schwameder used a model with 18 muscles in each leg, compared to eight in the current study, which could explain the higher forces in the iliopsoas, hamstrings, vastus and soleus than in [43]. The force in the gluteals was lower, and the force in the rectus femoris, gastrocnemius and tibialis anterior was similar to [43].

The relative contribution of the different joints to the total metabolic cost in the level trial at 1.3 m/s was smallest for the knee, between 13% and 26%, and similar between the hip and ankle, with the hip slightly smaller (between 29% and 41%) than the ankle (34% and 49%). Notably, model KIMR15 had the most unequal distribution between the hip and ankle, with 29% and 49%, while the other models had a difference in metabolic cost between these joints of less than 12%. These contributions are similar to contributions of different joints in guinea fowl walking (24%, 37%, and 38% for the knee, hip and ankle, respectively), measured using their blood flow [44]. In [26], the relative contribution was reported for tracking simulations. For these simulation, the energy was distributed more evenly between all three joints, with the knee contributing between 27% and 34%, the hip between 23% and 39%, and the ankle between 29% and 50%, while the absolute metabolic cost varied greatly between models [26]. Therefore, when creating simulation of walking, some energy minimization is required to distribute energy between joints more accurately.

Similar trends existed between the metabolic energy models in the breakdown of the metabolic cost per joint, though differences exist in the breakdown for the specific trials (see Fig 5). Since no measurements were taken, it cannot be said which model is more correct. However, the trends with speed and slope were very similar between models. The hip was mainly responsible for the change of energy expended with the slopes. When walking uphill, the energy expended in the hip increased, while it decreased when walking downhill. Between the speeds, the most obvious difference was an increase in energy expended in the ankle for the larger speed, while less energy is expended in the knee, such that the total metabolic cost remained similar. More energy was expended in the knee at 0.8 m/s, while more energy was expended in the ankle at 1.3 m/s.

Fig 2 showed a match with previous work [26]. The results for models MINE97, BHAR04, HOUD06, LICH05, and UMBE03 were very similar to Fig 2 in [26]. Note that model UMBE03 in [26] does not allow negative work, so the result differs for lengthening.

Commonly, muscle activations are found in gait analysis by static optimization [45] or computed muscle control (CMC) [46]. Static optimization only finds muscle activations, whereas model BHAR04 and model UMBE03 require activation and stimulation. All models, except model KIMR15, require the contractile element length and velocity. CMC requires a full-body model and markerset to solve for these variables. The approach in the current study required only a lower-extremity model and six markers. A larger markerset was used to aid data processing in Cortex. Similar to static optimization [45], solutions were robust to changes in the objective function (Eq. 9). Optimizations with objectives of cubed activation, with and without muscle volume weighting, yielded similar muscle forces.

Other metabolic models were recently developed by Uchida et al. [22] and by Tsianos et al. [25]. The model by Uchida et al. was very similar to model UMBE03, but it used a different method to determine the amount of fast twitch and slow twitch fibers. The results were very similar to model UMBE03, so they were not reported separately. The model by Tsianos et al. was not compatible with the musculoskeletal model that was used, since it is created for a different muscle model than a Hill-type muscle model, and was therefore not implemented.

Several equations have been developed to determine the energetic equivalent of pulmonary gas exchange measurements, for which an overview is given by Kipp et al. [47]. The analysis was repeated with the equation presented by Peronnet [48]. The repeated measures correlation was the same using Peronnet’s equation as when Weir’s equation was used, while the slopes were between 0.03 and 0.08 larger. However, the RMS errors were higher, except for the downhill and level trials of model MARG68 and MINE97. It was also found in the sensitivity analysis that the RMS error was affected by the isometric muscle force. Therefore, no further conclusions were drawn for the absolute predictions of the models.

The result of the current study are dependent not only on the metabolic energy models, but also on the methods used to find the inputs (e.g. muscle states) to these models and the measured metabolic cost that was used for validation. Fig 3 showed that the pattern of the muscle forces in the level trial at 1.3 m/s was similar to EMG data, while a comparison with [43] showed similar trends with slopes, which supports that the input to the models was accurate. The measured metabolic cost was found by subtracting resting metabolic cost that was found by standing. However, a bias could be present, since the calculated metabolic cost might not exactly represent this measured cost. Additionally, a two dimensional model of the lower part of the body was used in the current study, while some metabolic cost is associated with arm swing, and control of non-sagittal rotations in the lower extremity. This could cause the underestimation of metabolic cost that was reported in the current study. However, the high correlations indicate that a two dimensional musculoskeletal model in combination with a metabolic cost model is applicable in studies where the absolute error is not important. This is supported by previous work that showed that EMG activity of the vastus and soleus can explain 96% of the variance in metabolic cost of inclined walking [49].

Subjects were not asked to refrain from eating before the experiment. It is known that the peak influence of food on resting metabolic rate occurs after about 60 minutes for young adults [50], after which it slowly disappears [51]. Since the experimental set-up took at least 90 minutes, after which the measurements were taken relatively fast, and in a different order for each subject, it is expected that the effect is small. Using data from [50], and assuming the worst-case scenario that measurements were taken between 90 and 150 minutes after food intake, the effect would be around 0.2 W/kg, which is similar to the standard deviation between subjects and the difference between Weir’s and Peronnet’s equation.

In summary, we have studied the ability of seven metabolic energy models to represent changes in energy cost of walking due to an altered environment. All models correlated well with the metabolic cost of walking measured with indirect calorimetry, with correlation coefficients of at least 0.9. The correlation of models BHAR04 and LICH05 were highest at 0.96. Model MARG68 was best able to predict the absolute change in metabolic cost following a change in the dynamics. The subject’s mass affected the calculated metabolic energy expenditure for all metabolic energy models. All models were able to predict the trend of increased metabolic cost from downhill to level to uphill walking, while the metabolic cost calculated with model MINE97 was most similar between the two speeds, as was also observed in the measured metabolic cost.

## Supporting information

**S1 File. Metabolic Energy Models.** A description of the metabolic energy models that were compared.

**S2 File. Muscle Model.** A description of the muscle model that was used to determine the stimulation, activation, and other muscle parameters.

**S3 File. Repeated Measures Model with Weight-based Intercept.** This supplementary file describes a repeated measures model with the intercept based on the subject’s weight. This model can be used to determine the absolute metabolic energy expenditure using a metabolic energy model more accurately.

**S1 Table. Marker description.** Description of markers that were used in the experiment.

**S1 Fig. Placement of markers on the body.** Illustration of the placement of the markers that were used in the experiment on the body.

## Acknowledgments

The authors would like to thank Dr. Ken Sparks and Chelsea Turowski for their help with gathering experimental data.

