## Supplemental File 1 for "Metabolic cost calculations of gait using musculoskeletal energy models, a comparison study"

### S1: Metabolic Energy Models

#### S1.1: Model BHAR04

Model BHAR04 calculates the metabolic rate in W as a sum of the activation heat rate, $\dot{h}_A$, the maintenance heat rate, $\dot{h}_M$, the shortening/lengthening heat rate, $\dot{h}_{SL}$, and the work, $w$ [1].

$$\dot{E} = \dot{h}_A + \dot{h}_M + \dot{h}_{SL} + w \tag{1}$$

The activation heat rate is determined as follows:

$$\dot{h}_A = \phi m_{mus}(A_f u_f f_{FT} + A_s u_s f_{ST}) \tag{2}$$

where $\phi$ is a decay function, $m_{mus}$ is the muscle mass, $A_f = 133$ W/kg and $A_s = 40$ W/kg are activation heat rate constant for fast twitch and slow twitch muscles, respectively, $u_f$ and $u_s$ are the stimulation levels of the fast twitch and slow twitch muscles, and $f_{FT}$ and $f_{ST}$ are the percentages of fast twitch and slow twitch muscles [1].

$\phi$ is determined as follows:

$$\phi = 0.06 + \exp\left(\frac{-t_{stim}u}{\tau_\phi}\right) \tag{3}$$

where $t_{stim}$ is the time the muscle is activated above 0.1, $u(t)$ is the muscle stimulation, and $\tau_\phi$ is the decay time constant [1].

The stimulation of the slow twitch and fast twitch muscles are determined as follows:

$$u_f = 1 - \cos\left(\frac{\pi}{2}u(t)\right) \tag{4}$$

$$u_s = \sin\left(\frac{\pi}{2}u(t)\right) \tag{5}$$

The maintenance heat rate is determined as follows:

$$\dot{h}_M = l_M m_{mus}(M_f u_f f_{FT} + M_s u_s f_{ST}) \tag{6}$$

where $l_M$ is the length dependence of the maintenance heat rate (see Fig. 2 in [1]), and $M_f = 111$ W/kg and $M_s = 74$ W/kg are maintenance heat rate constants [1].

The shortening-lengthening heat rate is determined as follows:

$$\dot{h}_{SL} = -\alpha v_{CE} \tag{7}$$

where $v_{CE}$ is the contractile element velocity, negative for shortening, and $\alpha$ is a constant that is different for shortening and lengthening:

$$\alpha = 0.16a(t)F_{max}f(l_{CE}(t)) + 0.18F_{CE}(t) \quad v_{CE} \leq 0 \tag{8}$$

$$\alpha = 0.157F_{CE} \quad v_{CE} > 0 \tag{9}$$

where $a(t)$ is the muscle activation, $F_{max}$ is the maximum isometric force, $f(l_{CE}(t))$ is the force-length relationship, and $F_{CE}(t)$ the current force in the muscle [1].

Finally, the work rate is determined as follows:

$$w = -v_{CE}F_{CE} \tag{10}$$

#### S1.2: Model HOUD06

Model HOUD06 [2] calculates the metabolic rate in W as a sum of the activation heat rate, $\dot{h}_A$, the maintenance heat rate, $\dot{h}_M$, the shortening/lengthening heat rate, $\dot{h}_{SL}$, and the work, $w$.

$$\dot{E} = \dot{h}_A + \dot{h}_M + \dot{h}_{SL} + w \tag{11}$$

**Table S1.1. Constants for short twitch and fast twitch fibers.**

| Variable | Fast twitch | Short twitch |
|---|---|---|
| $h_A$ | 52.5 W/kg | 10.98 W/kg |
| $\bar{h}_M$ | 97.5 W/kg | 13.42 W/kg |
| $k_1$ | 12 | 6 |
| $k_2$ | 14 | 8 |
| $\bar{h}_{SL}$ | 0.28 $F_{max}$ | 0.16 $F_{max}$ |

The activation heat rate is determined as follows:

$$\dot{h}_A = m_{mus}\bar{h}_A\nu\frac{1-\exp(-0.25-\frac{18.2}{\nu\nu_{max}})}{1-\exp(-0.25-\frac{18.2}{\nu_{max}})} \tag{12}$$

where $\bar{h}_A$ is the activation heat rate constant, $\nu = a(t)^2$ is the relative stimulation frequency, and $\nu_{max} = k_1 + k_2 a(t)$ is the maximum stimulation frequency. These constants are defined for short twitch and fast twitch fibers and calculated for each muscles by taking the product of the constant for short twitch fibers and the percentage of short twitch fibers, and adding to this the product of the constant for fast twitch fibers and the percentage of fast twitch fibers. These values are given in table S1.1 [2].

The maintenance heat rate is determined as follows:

$$\dot{h}_M = m_{mus}(\bar{h}_A + \bar{h}_M)a(t)\left(f(l_{CE}(t)) - \frac{\bar{h}_A}{\bar{h}_A + \bar{h}_M}\right) \tag{13}$$

where $\bar{h}_M$ is the maintenance heat rate constant, different for short twitch and fast twitch fibers, as given in table S1.1 [2].

The shortening-lengthening heat rate is calculated as follows:

$$\dot{h}_{SL} = \bar{h}_{SL}a(t)f(l_{CE}(t)) - v_{CE} \qquad \text{where: } v_{CE} < 0 \tag{14}$$

where $\bar{h}_{SL}$ is given in table S1.1 for short twitch and fast twitch fibers [2].

#### S1.3: Model UMBE03

Model UMBE03 determines the metabolic rate per muscle in W as follows:

if $l_{CE} \leq l_{CE(OPT)}$

$$\dot{E} = m_{mus}(h_{AM}A_{AM}S + h_{SL}S) - w \tag{15}$$

if $l_{CE} > l_{CE(OPT)}$

$$\dot{E} = m_{mus}((0.4h_{AM} + 0.6h_{AM}f(l_{CE}))A_{AM}S + h_{SL}Sf(l_{CE})) - w$$

where $\dot{h}_{AM}$ is the activation-maintenance heat rate, $A_{AM} = A^{0.6}$, and $S$ is a scaling factor, equal to 1.5 in aerobic conditions. When the fiber length is longer than optimal, $\dot{h}_{AM}$ is split up into two parts, 40% represents the activation heat rate, while 60% represents the activation heat rate, which is dependent on the location on the force-length relationship [3].

The following equation is used to find the activation-maintenance heat rate [3]:

$$\dot{h}_{AM} = 128f_{FT} + 25 \tag{16}$$

The shortening-lengthening heat rate, $\dot{h}_{SL}$ is different for shortening and lengthening velocity and is determined as follows:

$$\dot{h}_{SL} = \begin{cases} \left(-\alpha_{S(ST)}\tilde{v}_{CE}(1-f_{FT}) - \alpha_{S(FT)}\tilde{v}_{CE}f_{FT}\right)A_s & \text{if } \tilde{v}_{CE} \leq 0 \\ \alpha_L\tilde{v}_{CE}A & \text{if } \tilde{v}_{CE} > 0 \end{cases} \tag{17}$$

where the first term, $\alpha_{S(ST)}\tilde{v}_{CE}(1 - f_{FT})$, cannot exceed 100 W/kg [3].

$\tilde{v}_{CE} = v_{CE}/l_{CE(OPT)}$ is the muscle fiber velocity in $s^{-1}$ normalized to the optimal fiber length, $l_{CE(OPT)}$. $\alpha_{S(ST)} = 100/\tilde{v}_{CE(MAX-ST)}$ and $\alpha_{S(FT)} = 153/\tilde{v}_{CE(MAX-FT)}$ are the shortening heat coefficients for slow twitch (ST) and FT fibers, respectively, in J/kg. They are dependent on the maximum fiber velocity of ST and FT fibers. $\alpha_L = 4\alpha_{S(ST)}$ is the lengthening heat coefficient, which is based on experimental data [3].

$A$ and $A_S$ are scaling factors that depend on the stimulation and activation:

$$A = \begin{cases} u & \text{when } u > a \\ (u + a)/2 & \text{when } u \leq a \end{cases} \tag{18}$$

$$A_S = \begin{cases} A^2 & \text{if } \tilde{v}_{CE} \leq 0 \\ A & \text{if } \tilde{v}_{CE} > 0 \end{cases} \tag{19}$$

#### S1.4: Model LICH05

Shortening velocity is positive in model LICH05 [4]. This model calculates the metabolic rate as a sum of the heat rate, $\dot{h}$, normalized to optimal fiber length and maximum isometric force, and the work:

$$\dot{E} = \dot{h} l_{CE(OPT)} F_{max} + w \tag{20}$$

The heat rate is a sum of the shortening-lengthening heat rate and the maintenance heat rate:

$$\dot{h} = (0.3a\dot{h}_M + 0.7af(l_{CE}(t))\dot{h}_M + af(l_{CE}(t))\dot{h}_{SL} \tag{21}$$

where 30% of the maintenance heat rate represents activation, which is only scaled by the activation [4].

The maintenance heat rate is determined as follows:

$$\dot{h}_M = \begin{cases} \gamma \frac{\tilde{v}_{CE(max)}}{G^2} & \text{if:} \quad v_{CE}(t) > 0 \\ 0.3\gamma \frac{\tilde{v}_{CE(max)}}{G^2} + 0.7\gamma \frac{\tilde{v}_{CE(max)}}{G^2} \exp(-7\tilde{v}_{CE(max)}(g(v_{CE}(t)) - 1)) & \text{if:} \quad v_{CE}(t) \leq 0 \end{cases} \tag{22}$$

where $\gamma$ is a heat rate value that decays with the stimulation time, $G = 4$ is the curvature of the force-velocity curve, and $g(v_{CE})$ is the location on the force-velocity relationship [4].

$\gamma$ is determined as follows [4]:

$$\gamma = 0.8 \exp(-0.72 t_{stim}) + 0.175 \exp(-0.022 t_{stim}) \tag{23}$$

The shortening-lengthening heat rate is determined as follows [4]:

$$\dot{h}_{SL} = \begin{cases} \frac{\tilde{v}_{CE}}{G} & \text{if:} \quad v_{CE}(t) > 0 \\ -0.5g(v_{CE}(t))v_{CE}(t) & \text{if:} \quad v_{CE}(t) \leq 0 \end{cases} \tag{24}$$
