## Supplemental Table 1 for "Metabolic cost calculations of gait using musculoskeletal energy models, a comparison study"

**Table S1: Marker description**

| No. | Name | Position |
|---|---|---|
| 1 | T10 | 10th thoracic vertebrae |
| 2 | SACR | Sacrum bone |
| 3 | NAVE | Navel |
| 4 | XYPh | Xyphoid process |
| 5 | STRN | Sternum |
| 6 | LASIS | Pelvic bone left front |
| 7 | RASIS | Pelvic bone right front |
| 8 | LPSIS | Pelvic bone left back |
| 9 | RPSIS | Pelvic bone right back |
| 10 | LGTRO | Left greater trochanter of femur |
| 11 | FLTHI | Left thigh |
| 12 | LLEK | Left lateral epicondyle of the knee |
| 13 | LATI | Left anterior of the tibia |
| 14 | LLM | Left lateral malleolus of the ankle |
| 15 | LHEE | Left heel |
| 16 | LTOE | Left toe |
| 17 | LMT5 | Left 5th metatarsal |
| 18 | RGTRO | Right trochanter major of the femur |
| 19 | FRTHI | Right thigh |
| 20 | RLEK | Right lateral epicondyle of the knee |
| 21 | RATI | Right anterior of the tibia |
| 22 | RLM | Right lateral malleolus of the ankle |
| 23 | RHEE | Right heel |
| 24 | RTOE | Right toe |
| 25 | RMT5 | Right 5th metatarsal |
| 26 | LSHO | Left Shoulder |
| 27 | RSHO | Right Shoulder |
