## Supplemental File 2 for "Metabolic cost calculations of gait using musculoskeletal energy models, a comparison study"

### S2: Muscle Model

Fig. S2.1 shows the three element Hill-type muscle that was used in this work. It consists of a contractile element, parallel elastic element and series elastic element. Its input is the stimulation, $u$, and the states are the activation, $a$, and the contractile element length, $l_{CE}$. The force in the contractile element is determined as follows:

$$F_{CE} = af(l_{CE})g(v_{CE})F_{max} \tag{1}$$

where $f(l_{CE})$ is the force-length relationship, and $g(v_{CE})$ is the force-velocity relationship.

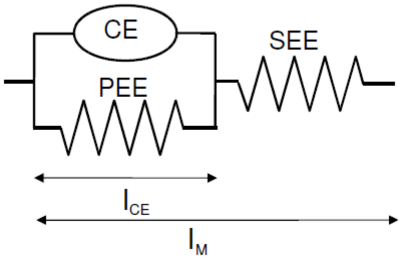


**Fig S2.1. Three element Hill-type muscle with contractile element, parallel elastic element, and series elastic element.**

The force-length relationship was as follows:

$$f(l_{CE}) = \exp\left(\frac{-(l_{CE} - l_{CE(OPT)})^2}{(Wl_{CE(OPT)})^2}\right) \tag{2}$$

The force-velocity equation was defined as follows:

$$g(v_{CE}) = \begin{cases} \frac{v_{CE(max)}+v_{CE}}{v_{CE(max)}-v_{CE}/A_{hill}} & \text{if} \quad v_{CE} < 0 \\ \frac{g_{max}v_{CE}+c}{v_{CE}+c} & \text{if} \quad v_{CE} \geq 0 \end{cases} \tag{3}$$

$$\text{with} \quad c = \frac{v_{CE(max)}A_{hill}(g_{max} - 1)}{A_{hill} + 1} \tag{4}$$

The parallel and series elastic elements are modeled as quadratic springs. Their force are found based on the model presented by McLean et al. [1]:

$$F(l) = \begin{cases} k_1(l - l_{slack}) & if l \leq l_{slack} \\ k_1(l - l_{slack}) + k_2(l - l_{slack})^2 & if l > l_{slack} \end{cases} \tag{5}$$

where $l$ denotes the length of the element and $l_{slack}$ the slack length. $k_1$ and $k_2$ are stiffness constants. $k_1 = 0.01\ F_{max}$/m represents a small linear stiffness, which was added to aid the optimization. It is equal to. $k_2$ is equal to the following:

$$k_2(PEE) = \frac{F_{max}k_{PEE}}{l^2_{CE(OPT)}} \tag{6}$$

$$k_2(SEE) = \frac{F_{max}}{(u_{max}l_{CE(OPT)})^2} \tag{7}$$

where $k_{PEE} = 1$ and $u_{max} = 0.04$ are dimensionless constants.

The muscle mass, $m_{mus}$ was determined using the maximum isometric force and the optimal fiber length:

$$m_{mus} = \frac{F_{max}}{\sigma}\rho l_{CE(OPT)} \tag{8}$$

where $\sigma = 25$ N/cm$^2$ is the muscle-specific stress [2] and $\rho = 1059.7$ kg/m$^3$ is the density of muscle.

Tab. S2.1 shows the maximum isometric force $F_{max}$, optimal fiber length, $l_{CE(OPT)}$, width of the force-length curve, the slack length of the parallel elastic element (PEE) and the series elastic element (SEE), the nominal muscle length, $l_m$, and the percentage of fast twitch fibers for each muscle. Several parameters parameters for each muscle. Several parameters were the same for each muscle, the activation time, $T_{act} = 0.01$ s, the deactivation time, $T_{deact} = 0.03$ s, the maximum shortening velocity, $v_{CE(max)} = 12\ l_{CE(OPT)}$/s, the maximum force during lengthening, $g_{max} = 1.5$ $F_{max}$, and the normalized hill constant, $A_{hill} = 0.25$.

**Table S2.1. Muscle Parameters**

| Muscle | $F_{max}[N]$ | $l_{CE(OPT)}$ [m] | Width | PEE slack $[l_{CE(OPT)}]$ | SEE slack [m] | $l_m$ [m] | % FT fibers |
|---|---|---|---|---|---|---|---|
| Iliopsoas | 1500 | 0.102 | 1.298 | 1.2 | 0.142 | 0.248 | 0.5 |
| Gluteals | 3000 | 0.2 | 0.625 | 1.2 | 0.157 | 0.271 | 0.45 |
| Hamstrings | 3000 | 0.104 | 1.197 | 1.2 | 0.334 | 0.383 | 0.35 |
| Rectus Femoris | 1200 | 0.081 | 1.443 | 1.4 | 0.398 | 0.474 | 0.65 |
| Vastus | 7000 | 0.093 | 0.627 | 1.4 | 0.223 | 0.271 | 0.5 |
| Gastrocnemius | 3000 | 0.055 | 1.039 | 1.2 | 0.42 | 0.487 | 0.5 |
| Soleus | 4000 | 0.055 | 1.039 | 1.2 | 0.245 | 0.284 | 0.2 |
| Tibialis Anterior | 2500 | 0.082 | 0.442 | 1.2 | 0.317 | 0.381 | 0.25 |
