## Supplementary figures and images for "Metabolic cost calculations of gait using musculoskeletal energy models, a comparison study"

### Supplemental Figure 1

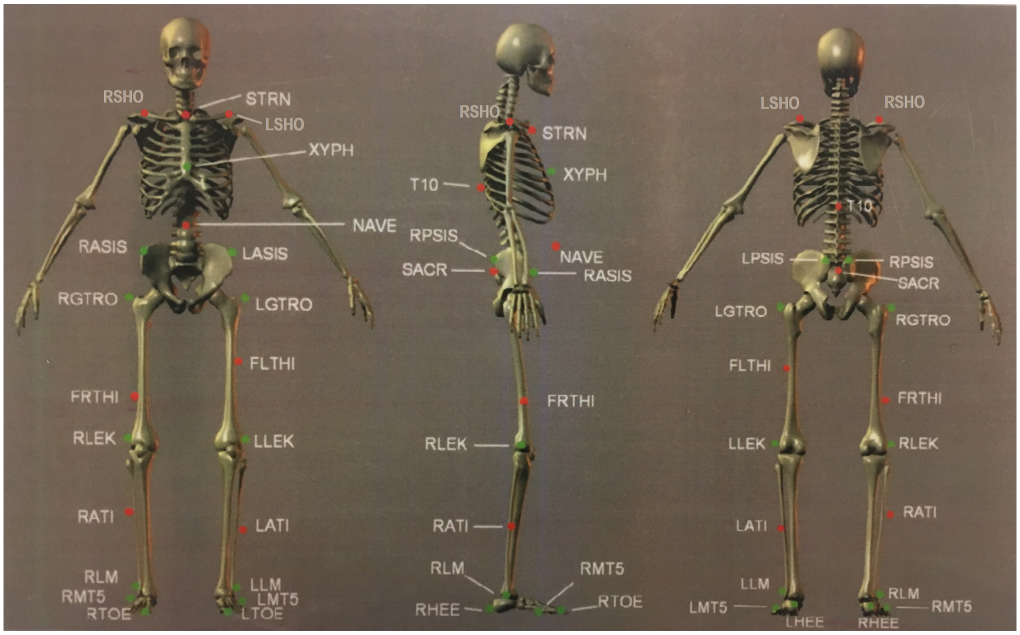

**Fig S1: Placement of markers on the body**
